# The adequacy of tissue microarrays in the assessment of inter- and intra-tumoural heterogeneity of infiltrating lymphocyte burden in leiomyosarcoma

**DOI:** 10.1101/412387

**Authors:** ATJ Lee, W Chew, MJ Smith, DC Strauss, C Fisher, AJ Hayes, I Judson, K Thway, RL Jones, PH Huang

**Author notes:** Correspondence: Dr. Paul Huang, Division of Molecular Pathology, The Institute of Cancer Research, London SW3 6JB, United Kingdom. These authors are joint senior authors (RL Jones, PH Huang).

## Abstract

The characterisation and clinical relevance of tumour-infiltrating lymphocytes (TILs) in leiomyosarcoma (LMS), a subtype of soft tissue sarcoma that exhibits histological heterogeneity, is not established. The use of tissue microarrays (TMA) in studies that profile TIL burden is attractive but given the potential for intra-tumoural heterogeneity to introduce sampling errors, the adequacy of this approach is undetermined. In this study, we assessed the histological inter-and intra-tumoural heterogeneity in TIL burden within a retrospective cohort of primary LMS specimens. Using a virtual TMA approach, we also analysed the optimal number of TMA cores required to provide an accurate representation of TIL burden in a full tissue section. We establish that LMS have generally low and spatially homogenous TIL burdens, although a small proportion exhibit higher levels and more heterogeneous distribution of TILs. We show that a conventional and practical number (1-3) of TMA cores is adequate for correct ordinal categorisation of tumours with high or low TIL burden, but that many more cores (≥ 11) is required to accurately estimate absolute TIL numbers. Our findings provide a benchmark for the design of future studies aiming to define the clinical relevance of the immune microenvironments of LMS and other sarcoma subtypes.

## Introduction

Tissue microarrays (TMAs) are useful diagnostic and research tools that permit high-throughput histological and molecular studies of up to several hundred tissue specimens simultaneously by arraying them into a paraffin block1. This approach offers several advantages over conventional examination of full tissue sections by minimising consumption of often limited tissue while providing various efficiencies in downstream sample processing and analysis. However, an inherent limitation to the use of TMAs is that, for each included specimen, only a small amount of tissue is sampled and arrayed, meaning that sampling error may lead to a distorted representation of the full tissue section. This limitation is of particular relevance in the study of tumour specimens, where intra-tumour spatial heterogeneity in terms of morphology and underlying molecular pathology is now well established in many cancer types^2^. Multiple studies have been undertaken to validate the TMA methodology in the assessment of various cancer biomarkers, with the aim of demonstrating that biomarker levels reported by TMAs are representative of results obtained when full sections are assessed. In this manner, it has been shown that expression levels of a diverse repertoire of tumour biomarkers are accurately reported through the assessment of TMAs, typically with the provision that between 1 and 3 replicate cores from each included tumour are assessed and aggregated^3–6^.

Investigating the immune microenvironment as a potential source of cancer biomarkers is an area of renewed research interest. It is now known that the presence or absence of immune-related factors such as tumour-infiltrating lymphocytes (TIL) can serve as powerful prognostic and/or predictive biomarkers across a range of cancer types^7–10^. TMAs continue to be frequently employed in studies that seek to investigate the potential role of TILs as putative biomarkers^7,11–15^. The success of such studies is dependent on the ability of the TMA approach to capture a sufficiently representative picture of the immune phenotypes present within the wider tumour. Observations of quantitative and qualitative spatial heterogeneity in the immune microenvironment of individual tumours of various cancer types call into question how well TMAs can provide such representation and whether the scope for sampling error renders them inappropriate for studies of tumour immunity^16–20^. There is currently little published evidence addressing this question^21–23^.

In this study, we assessed a cohort of leiomyosarcoma (LMS) tumour specimens to investigate the extent of inter-and intra-tumoural heterogeneity of TIL burden and how accurately TIL burden are represented by the TMA methodology, compared to full tumour sections. LMS are tumours of smooth muscle lineage and are one of the more common soft tissue sarcoma (STS) subtypes, representing 10-20% of all STS^24^. As with other STS subtypes, the immune microenvironment and its potential prognostic value is not well characterised in LMS. There is accumulating evidence that LMS is a disease that harbours extensive inter-and intra-tumoural genetic and morphological heterogeneity. For instance, recent genomic profiling analyses demonstrate that LMS is characterised by inter-tumour variability in somatic copy number alterations, a molecular characteristic found to have negative correlation with active anti-tumour immune response in a number of other cancers^25,26^. Furthermore, clinical evidence suggest that a small minority of LMS respond to immune checkpoint inhibitor therapy^27–29^. As such, inter-patient differences in the immune microenvironment of LMS may be useful for predicting response to therapy and prognosis. LMS often present with large primary tumours that exhibit intra-tumoural morphological heterogeneity of tumour cells and associated stroma, and consequently may also display intra-tumoural TIL heterogeneity^30^. To assess the suitability of TMAs for profiling TIL burden in LMS, we sought to address two questions in this study: 1) What is the extent of inter-and intra-tumour heterogeneity of TIL burden in LMS, by comparing related tumour blocks from spatially distinct areas of primary tumours and 2) how many TMA cores are required to provide sufficient representation of the TIL burden of the full tissue section?

## Materials and Methods

### Tumour sample selection and processing

Surgical resection specimens of primary LMS (n=47) and accompanying annotation of baseline clinicopathological variables were identified and retrieved through retrospective review of departmental database and medical notes at a single specialist cancer centre. Histological diagnosis was confirmed by a specialist sarcoma histopathologist (CF, KT). Where available, 5 blocks containing formalin-fixed paraffin-embedded (FFPE) viable tumour from spatially distinct areas (at least 2 blocks each from tumour margins and core) of the same primary tumour were selected. Newly prepared haematoxylin and eosin (H&E) slides from each block were assessed to confirm presence of viable tumour material. Immunohistochemical (IHC) staining of T lymphocyte markers (anti-CD3 [clone M0452, DAKO, 1:600 dilution], anti-CD4 [4B12, DAKO, 1:80] and anti-CD8 [C8/144B, DAKO, 1:100]) was performed on consecutive 4μm sections from each block **(See supplemental methods for further details)**. IHC staining for B lymphocytes (anti-CD20 [L26,DAKO, 1:400]) was performed on all blocks from an initial set of 19 tumours – this was not expanded to all tumours due to uniformly low numbers of infiltrating B cells in this initial set.

### IHC scoring

#### Full tissue sections

The number of CD3, CD4, CD8 and CD20 IHC-positive lymphocytes in ten non-adjacent, tumour-containing high-power fields (HPF) (x400 magnification, approx. area per HPF 0.31mm^2^) was manually counted by direct brightfield microscopy for each stained slide.

#### Virtual TMA (vTMA)

Digital microscopy images for slides stained for H&E, CD3 and CD8 from a single block from each of 47 cases were captured at x40 resolution using Nanozoomer-XR (Hamamatsu Photonics). 20 × 1mm diameter circular areas were randomly selected from viable-tumour areas on each H&E image. The corresponding areas were then selected on CD3 and CD8 digital slide images. Images of these areas at x10 magnification were exported as .tif files that were cropped to uniform 0.785mm^2^ circular areas in Image J^31^. Positive-staining TILs in these images were counted using ‘Particle analysis’ function of image J following optimization of pixel intensity, particle size and circularity thresholds – the selected configuration was associated with a bias of -0.52 cells, with 95% limits of agreement at -17 to +16 cells, as assessed by Bland-Altman analysis. Due to the presence of pleomorphic, CD4-staining histiocytes, we were unable to use this approach for counting of CD4+ TILs, and so this marker was not included in the vTMA experiment.

#### Physical TMA (pTMA)

Triplicate 1mm diameter cores were sampled from areas of viable tumour within donor blocks from 44/47 LMS and re-embedded in an arrayed recipient paraffin block. Consecutive 4μm sections from the arrayed block were stained for H&E, CD3 and CD8. After assessment of H&E slides to confirm viable tumour content, all CD3+ and CD8+ TILs were counted under direct brightfield microscopy. Average TIL number per 1mm core (referred to herein as ‘TIL/core’) was calculated from triplicate cores for each tumour.

### Statistical analysis

#### Degree of infiltrating lymphocyte burden across LMS cohort

To assess the extent of TIL burden in each of 47 LMS cases, an average number of infiltrating CD3+, CD4+ and CD8+ TIL per HPF (referred to herein as ‘TIL/HPF’) was calculated from 50 HPF per tumour (10 HPF from each of 5 related tumour blocks). Comparison of TIL burden of tumours from different anatomical sites of origin was performed using 1-way ANOVA of Log2-transformed average TIL/HPF values with Prism v7.0 (GraphPad Software Inc).

#### Inter-vs intra-tumour variance in TILs

To assess the variability in TIL burden between different blocks from the same surgical specimen, we assessed the relative contribution of inter-block variation (block effect) and inter-tumour variation (tumour effect) on the total amount of variance in TIL numbers within the 47 LMS cohort by (i) Log2 transformation of all raw TIL/HPF count values (ii) calculation of average TIL/HPF with 95% confidence interval for each tumour block (average of 10 HPF,) and across all 5 related blocks from each primary tumour (average of 50 HPF), and (iii) Ordinary 2 way ANOVA (Prism v7.0) to assess the percentage of total variability attributable to block effect, tumour effect, interaction between the two effects and residual variation.

#### Virtual TMA assessment of optimal core number

Automated counts of infiltrating CD3+ and CD8+ TIL in all 20 × 1mm vTMA cores for each tumour were used to calculate average TIL/core – this value was taken as representing the ‘true’ TIL burden of each tumour. Estimates of these true TIL burdens were then derived from the average TIL/core from all possible combinations of between 2 and 19 randomly-selected cores. The percentage of estimates generated from *n* cores that fell within the following prescribed boundaries were then calculated: a) within +/- 20% of true TIL burden; b) within correct (i.e. same as true TIL burden) side of dichotomised ‘high/low’ boundary set at median of true TIL burdens from 47 LMS cohort; c) within correct quartile of 47 LMS cohort.

#### Assessment of accuracy of triplicate cores within a physical TMA

The differences between Log2-transformed values of the estimated average TIL/core values derived from the pTMA and the true TIL burdens derived from the vTMA were calculated and plotted against the average of the two values in a Bland Altman plot along with 95% levels of agreement (Prism v7.0).

Average of TIL/core values derived from triplicate cores within pTMA were used to identify each included LMS as having a ‘high’ or ‘low’ TIL burden, relative to the cohort median of true TIL burdens, as defined in the vTMA experiment. This high/low identification was then compared to a ‘gold standard’ high/low allocation, defined as the ‘true’ TIL burden of that tumour as derived from all 20 vTMA cores. Accuracy (%) of pTMA was defined as 100*(True Positive + True Negative)/ (True Positive+ False Positive + True Negative+ False Negative)

### Research ethics

Use of archival FFPE tumour samples and linked anonymised patient was approved by Institutional Review Board as part of the PROSPECTUS study, a Royal Marsden-sponsored non-interventional translational protocol (CCR 4371, REC 16/EE/0213).

## Results

### Patient and tumour characteristics

Adequate tumour material was identified for 47 patients with a confirmed diagnosis of LMS who had undergone radical resection of primary tumour (baseline clinicopathological variables are summarised in **Table 1**). A majority of tumours were >5cm in maximal dimension, and 44% >10cm. Six tumours (13%) were low grade.

**Table 1.** Baseline clinicopathological status of 47 patients with primary LMS. RP=retroperitoneal. AP = abdominopelvic.

|  | **Overall** | **Intra-cavity**<br>**(exc. Uterine)** | **Uterine** | **Extremity/**<br>**Trunk** |
| --- | --- | --- | --- | --- |
| **N (%)** | 47 | 27 (57%) | 8 (17%) | 12 (26%) |
| **Anatomical**<br>**position** | - | RP - 20 (74%)<br>AP- 7 (26%) | - | Lower limb - 6 (50%)<br>Upper limb - 3 (25%)<br>Trunk - 3 (25%) |
| **Average**<br>**age/years**<br>**(range)** | 61.5<br>(29.4 - 87.5) | 62.2<br>(30.6-87.0) | 46.4<br>(29.4-73.4) | 69.8<br>(53.2-87.5) |
| **% M:F** | 51:49 | 48:52 | 0:100 | 83:17 |
| **Max tumour**<br>**dimension:** |  |  |  |  |
| **<5cm** | 5 (11%) | 2 (7%) | 1 (13%) | 1 (8%) |
| **5-10cm** | 21 (45%) | 12 (44%) | 3 (38%) | 6 (50%) |
| **10-15cm** | 10 (21%) | 6 (22%) | 1 (13%) | 3 (25%) |
| **>15cm** | 11 (23%) | 7 (26%) | 3 (38%) | 1 (8%) |
| **Histological**<br>**grade:** |  |  |  |  |
| **1** | 6 (13%) | 6 (22%) | 0 | 0 |
| **2** | 25 (53%) | 16 (59%) | 3 (38%) | 6 (50%) |
| **3** | 16 (34%) | 5 (19%) | 5 (62%) | 6 (50%) |

### LMS are variably infiltrated by T lymphocytes

For each case in the cohort, 5 tissue blocks that sampled spatially distinct tumour areas were assessed for TIL burden (outlined in workflow in **Figure 1A**). IHC staining for CD3 was used as a global T lymphocyte marker, with staining of consecutive slides for CD8 and CD4 used as markers for cytotoxic and helper T cell subpopulations respectively. CD20 expression was used as a global marker for B lymphocytes. Positive-staining TILs in 10 non-adjacent HPF were counted in sections from each of 5 blocks per tumour, with the average of all 50 related HPF (equating to a total area of 15.5mm^2^ of assessed tumour) taken to represent the overall tumour TIL burden. Exemplar IHC images showing different degrees of CD3+ lymphocyte infiltration are shown in **Figure 1B**. The distribution of overall tumour TIL burdens for each lymphocyte marker across the cohort is shown in **Figure 1C**. The cohort medians of average TIL/HPF were CD3: 16.5 (IQR 11.3-30.9), CD4: 10.5 (IQR 5.5-18.9), CD8: 16.1 (IQR 7.2-23.0). These median values are below the ‘low infiltration’ thresholds currently used in studies of TILs in other well-studied cancer types such as melanoma, non-small cell lung cancer (NSCLC) and colorectal cancer^33–37^.

**Figure 1.**
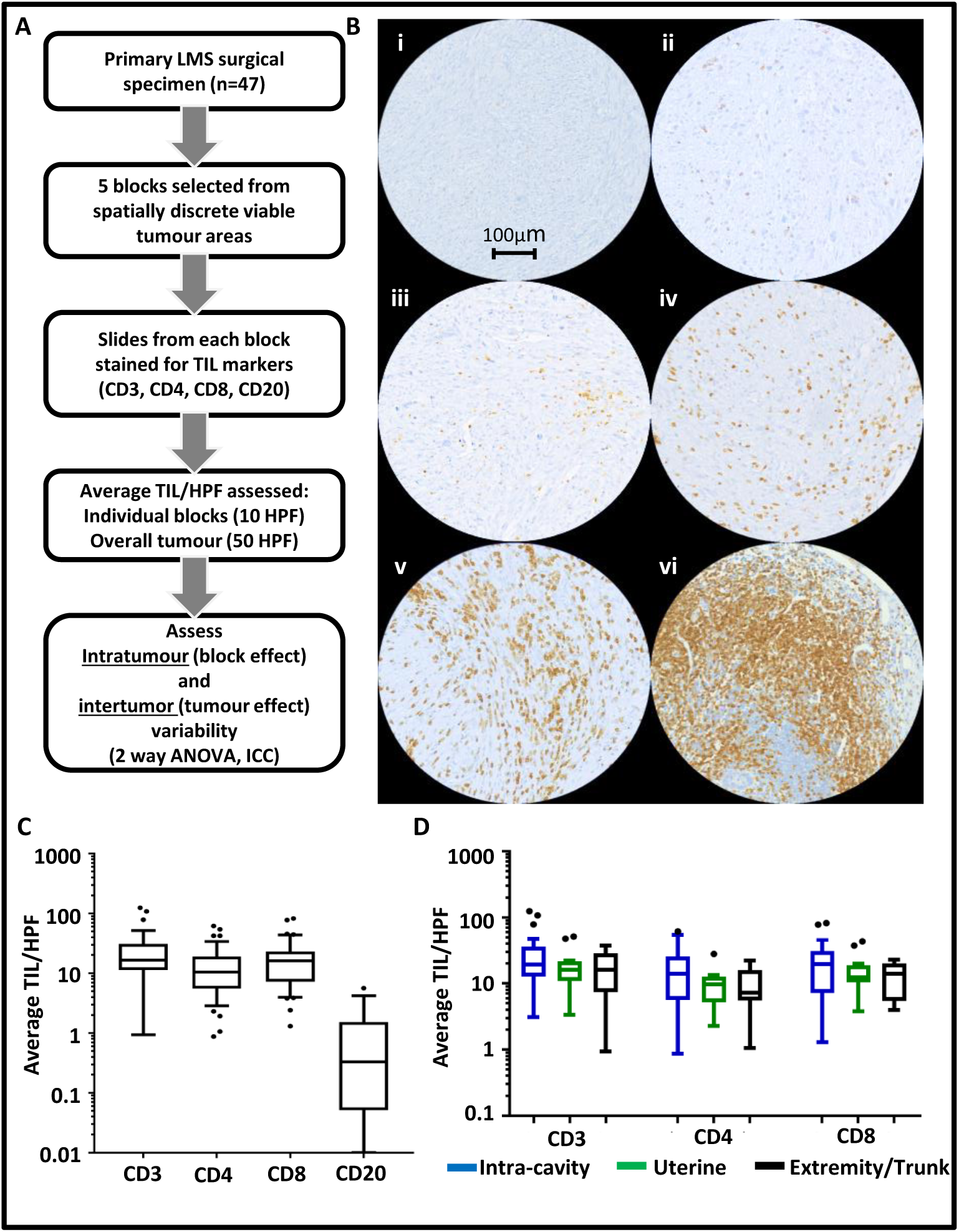
Evaluation of infiltrating T and B lymphocyte burden in LMS. **A.** Workflow of experimental approach. **B.** Representative areas from different CD3-stained LMS demonstrating range of infiltrating CD3+ T lymphocyte burdens. Densities are (i) 1, (ii) 35, (iii) 100, (iv) 250 and (v) 800 TIL/HPF (vi) positive control tissue (appendix) with 1400 TIL/HPF. **C.** Tukey box and tail plots showing overall lymphocyte burdens (average number of tumour-infiltrating lymphocytes (TIL) per x400 high-powered fields (HPF), calculated from 50 HPF) in LMS cohort based on IHC staining for CD3, CD4, CD8 (n=47) and CD20 (n=19) of LMS **D.** Tukey box and tail plots showing distribution of CD3+, CD4+ and CD8+ TIL burdens within 47 LMS cohort when stratified by site of tumour origin. 1 way ANOVA of Log2-transformed values demonstrates no significant differences in T lymphocyte counts between tumours of different site of origin

A median CD20+ TIL/HPF of 0.3 (0.1 – 1.5), indicated the near-absence of infiltrating B cells in a subset of 19 tumours. Across the different T lymphocyte markers, a dynamic range of 2-3 orders of magnitude (e.g. CD3 range 1-124 TIL/HPF) was seen in the extent of TIL burden between individual tumours **(Figure 1B)**. No significant differences in T lymphocyte burden was seen when comparing LMS from different anatomical sites of origin **(Figure 1D)**. These data indicate that marked variation in TIL burden is seen among individual LMS cases in a manner that was not associated with anatomical site of origin, and that LMS generally have a lower TIL burden than other, well-studied epithelial tumour types.

### Inter-tumour heterogeneity in TIL burden of LMS greatly outweighs intra-tumour heterogeneity

Having established that overall TIL burdens can vary between individual LMS tumours, we assessed the extent of heterogeneity in average TIL/HPF between blocks taken from different regions from the same LMS specimen **(Figure 1A)**.

Average TIL burden (stated as TIL/HPF) from each of 5 sampled blocks from the 47 cases are shown aligned with overall tumour average values in **Figure 2A**. These data demonstrate that in most LMS cases, all the blocks from the same tumour had similar TIL/HPF values, suggesting low levels of intra-tumoural heterogeneity in these cases. However, in a subset of 10 cases (21%), TIL/HPF values varied widely between individual blocks from the same tumour, indicating higher levels of heterogeneity in TIL distribution. Differences in the extent of intra-tumoural TIL heterogeneity between individual LMS tumours is further exemplified in 3 cases, as illustrated in **Figure 2B.** Notably, the tumours with the greatest extent of intra-tumour TIL heterogeneity tended to be those cases with the highest overall TIL burdens **(Figure 2A)**.

**Figure 2.**
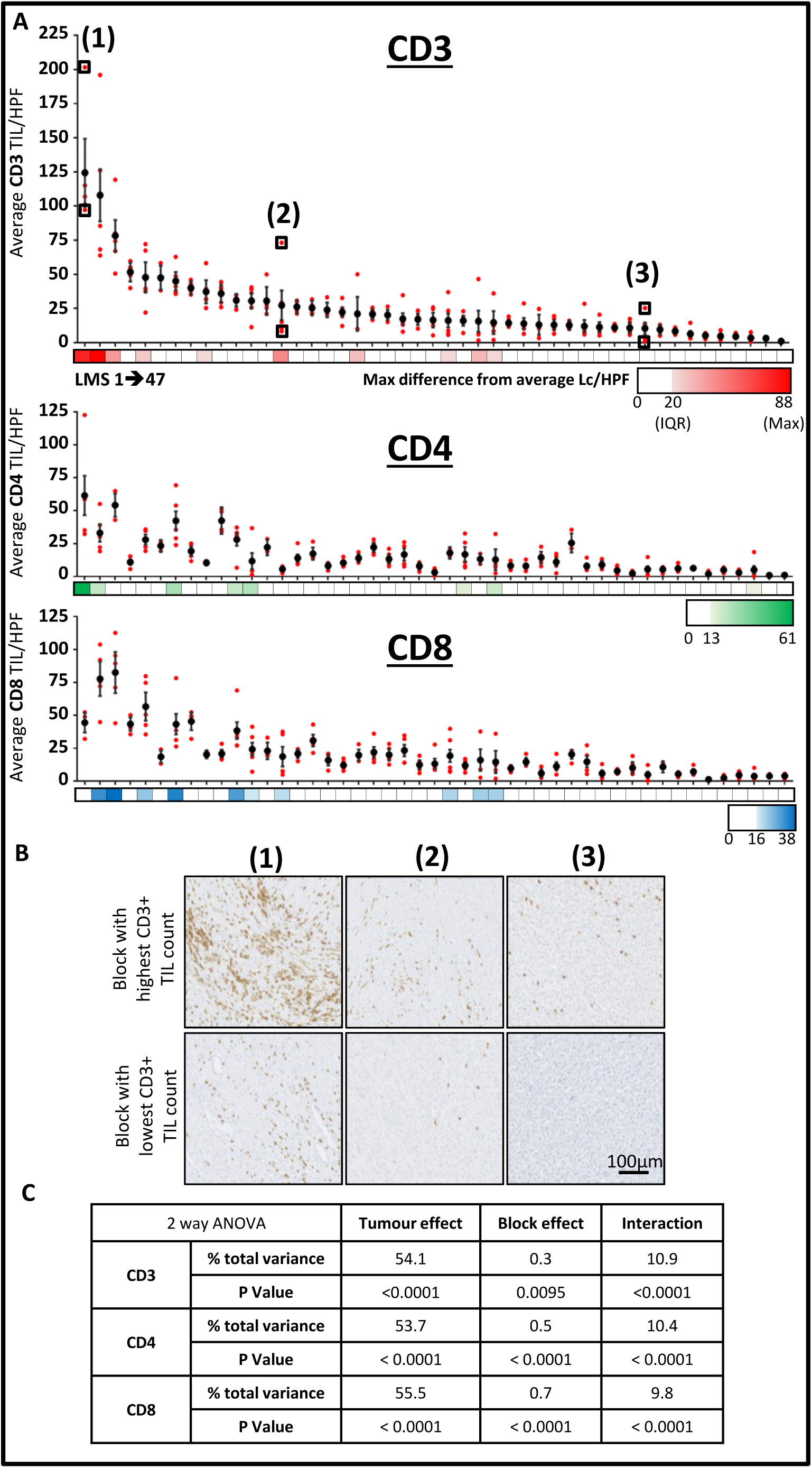
Assessment of inter-and intra-tumour heterogeneity of TIL burden in LMS. **A.** Dot plot shows average CD3+, CD4+ and CD8+ TIL/HPF values for each of 47 LMS tumours (vertically aligned), with overall tumour value (+/-95% confidence interval) and individual constituent blocks values shown with in black and red respectively. Colour bars demonstrate maximum difference of any related tumour block from overall tumour average, with zero, cohort interquartile range (IQR), and maximum difference values shown on colour key for each lymphocyte marker. **B.** Representative IHC images at x40 magnification demonstrate CD3+ TIL burden between the most and least densely infiltrated blocks from three tumours as indicated in (**A). C.** Table summarising results from three separate 2-way ANOVA analyses that identifies the contribution of intra-tumour (block effect) and inter-tumour (tumour effect) variance to the overall total amount of variance in lymphocyte counts for CD3, CD4 and CD8 within the 47 LMS cohort.

We performed 2 way ANOVA to objectively assess the relative extent that intra-tumoural heterogeneity (i.e. variation between blocks from the same tumour – ‘block effect’) and inter-tumoural heterogeneity (i.e. variation in overall TIL burdens between different tumours – ‘tumour effect’) contributed to the overall amount of variation in TIL burden within the cohort **(Figure 2C)**. We found that block effect had a much smaller contribution to the overall amount of variance compared to the contribution of tumour effect between cases within the cohort. Tumour effect accounted for 54.1%, 53.7% and 55.5% of total variance in lymphocyte counts for CD3, CD4 and CD8 respectively, while block effect contributed to only 0.3%, 0.5% and 0.7% total variance for the same respective markers. Significant interaction between tumour and block effect was detected for all three T lymphocyte measurements, in keeping with the observation that a greater degree of intra-tumour variance is observed in tumours with higher TIL burdens.

Taken together, these results indicate that while intra-tumoural heterogeneity was observed in a subset of LMS cases with higher overall TIL levels, intra-tumoural heterogeneity in TIL burden across the cohort was outweighed by the extent of inter-tumoural heterogeneity.

### Optimal number of cores to ensure representativeness of tissue microarrays depends on required degree of accuracy

To address the question of how many TMA cores must be sampled from a tumour to provide adequate representation of the overall TIL burden of a tumour, we devised an *in silico* ‘virtual TMA’ (vTMA) that would allow for the iterative sampling of a number of cores that would be impractical for a physical TMA. We then assessed how many cores were required to produce an estimate of TIL burden that either (i) accurately recapitulated the true TIL burden of a tumour, or (ii) was sufficiently accurate to identify whether a tumour had high or low TIL burden, relative to the median or quartile TIL values of the entire cohort– this second approach was based on the observation that, in many published studies that have demonstrated clinical relevance of TIL numbers, similar rank-based categorisation was used, often based on dichotomisation around cohort median value^37^.

For each of 47 LMS cases, digital microscopy images were taken of H&E, CD3 and CD8 stained whole sections of a single tumour block. 20 × 1mm circular ‘core’ areas (total area. 15.7mm^2^ - equivalent to approximately 50 HPF) were selected on H&E images, with the number of TILs within the corresponding areas (TIL/core) on CD3 and CD8 stained slides digitally counted **(Figure 3A)**. For each tumour, the average TIL/core from each of every possible combination of 2 out of 20 cores, 3 out of 20 cores, and so on, were calculated **(Figure 3B).** The average TIL/core of all 20 cores was taken to represent the ‘true’ overall TIL burden of each tumour. For each of the 47 tumours, we assessed how many cores needed to be sampled in order for >80% of possible combinations to produce an estimated TIL burden that fell within each of three different thresholds: (i)+/-20% of true TIL burden, (ii) same side of cohort median or (iii) in same cohort quartile as true TIL burden across the entire cohort (**Figure 3C)**.

**Figure 3.**
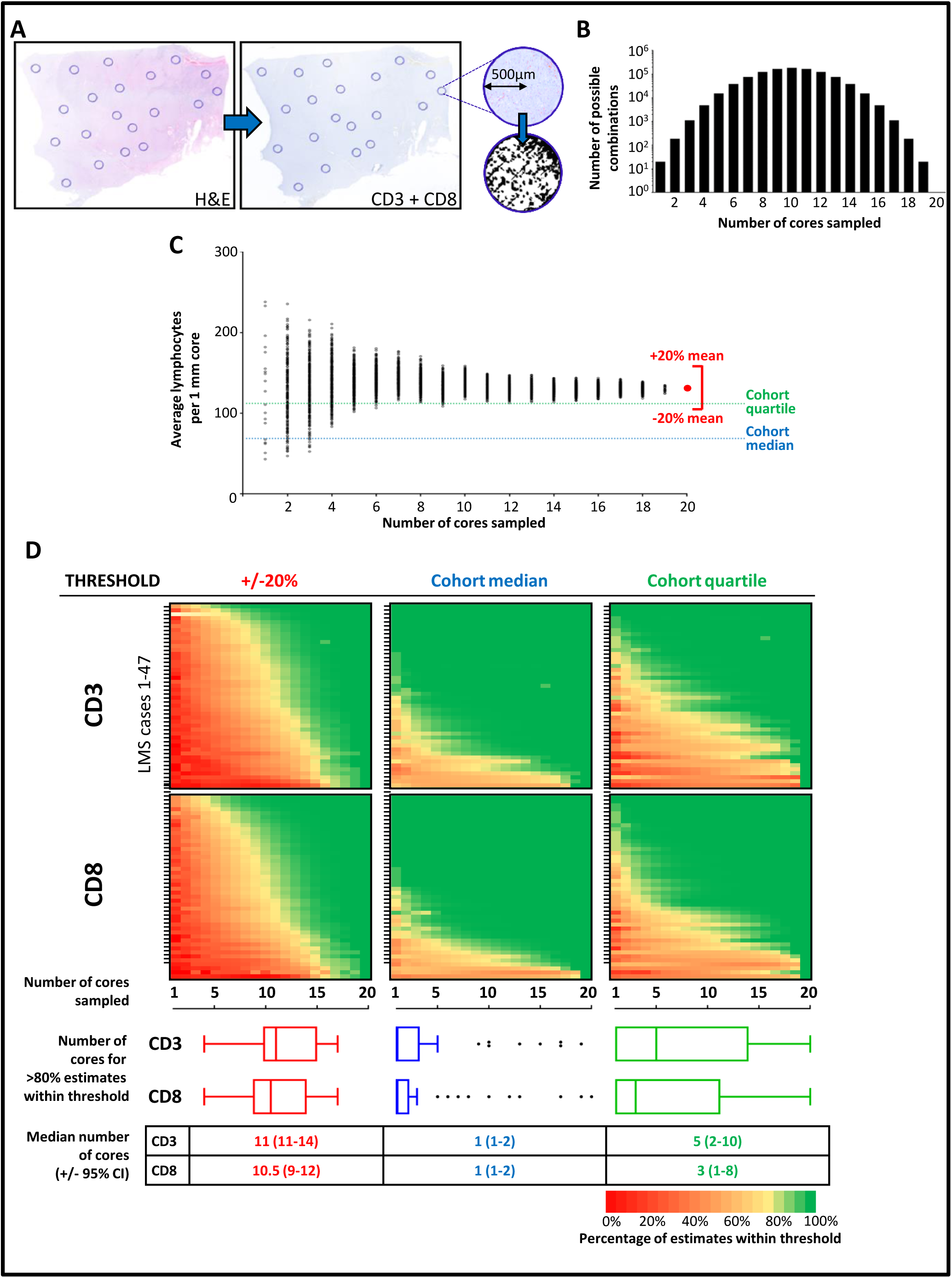
Optimal number of TMA cores relates to required degree of accuracy for assessment of lymphocyte infiltration. **A.** Overview diagram of process for selection of virtual TMA cores and T lymphocyte counting. For each of 47 LMS, a digital H&E slide from a representative block was marked for 20 × 1mm diameter areas, encompassing spatial and any morphological heterogeneity with section. Selected core areas were mapped on to corresponding CD3 and CD8-stained sections. Core areas were isolated as individual digital images. Number of IHC-positive lymphocytes in each core area was digitally counted. **B.** Bar chart showing number of possible combination of cores when between 2-20 cores are assessed. Average lymphocyte count per core (TIL/core) was calculated for all possible combinations for each tumour. **C.** Dot plot showing all possible average lymphocyte counts (number indicated in **B**) for a single exemplar tumour when 1-20 cores are selected. Average of all 20 cores (red dot) taken to be represent overall TIL burden for that tumour. For each tumour, the number of cores that needed to be sampled in order for >80% calculated averages to fall within either (i) +/-20% of ‘true TIL burden’ for corresponding tumour, (ii) correct side of cohort median TIL value (CD3 median = 69 TIL/core; CD8 median = 59 TIL/core), or (iii) within correct cohort quartile (CD3 IQR = 18-110 TIL/core; CD8 IQR = 19-121 TIL/core). In this illustrated exemplar case, overall CD3+ TIL burden is above 3rd quartile (Q75 = 110). **D**. Colour plots indicating percentage of systematically calculated average lymphocyte counts from all possible combinations of between 1-20 cores to fall within stated threshold (+/-20%, cohort median or cohort quartile) Tukey box and tail plots indicate cohort distribution of number of cores required for >80% of estimates to fall within stated threshold. Table summarises cohort median number of cores required >80% of estimates to fall within stated threshold (+/- approx. 95% confidence interval).

A median of 11 cores (CD3 range 4-16, CD8 range 4-17) was required for >80% of estimated TIL burdens to fall within 20% of the ‘true’ CD3+ or CD8+ TIL burden for the corresponding tumour **(Figure 3D)**. However, for the majority of cases, only 1 core was required for >80% of estimated CD3+ or CD8+ TIL burdens to fall the same side of the cohort median as the corresponding true TIL burden. Similarly, a lower number of cores (median of 5 and 3 cores for CD3 and CD8 respectively) were required for >80% of estimated TIL burdens to fall in the same cohort quartile as the corresponding ‘true’ TIL burden. A minority of tumours required a greater number of cores for >80% of estimated TIL burdens to fall on the correct side of cohort median (8/47 and 6/47 requiring ≥8 cores for CD3 and CD8 respectively), primarily due to these tumours having true TIL burdens that lay close to median cut-off values (**Figure 3D**).

Taken together, these data indicate that a large and likely impractical number of TMA cores (11 cores) must be sampled in order to accurately recapitulate the true burden of infiltrating T lymphocytes in LMS. However, many studies that have described an association between TILs and clinical outcome ultimately applied cut-off thresholds to assign TIL counts into ordinal categories (e.g. ‘high’ or ‘low’ infiltration) that reflect relative rather than absolute degree of infiltration37. We found that sampling only 1 core was sufficient to correctly identify a majority of tumours as has having a ‘high’ or ‘low’ degree of infiltration, while 2-5 cores was adequate to correctly identify a majority of tumours as having ‘very low’, ‘low’, ‘high’ or ‘very high’ degree of TIL infiltration, based on categorical cut-offs at cohort quartiles. These data demonstrate that TMAs that employ a conventional number of cores (i.e. 1-3) would be sufficiently representative for studies where ordinal categorisation of TIL burden is planned. However, should precise quantification of the absolute value of true TIL burden be desired, a conventional TMA approach is not likely to be representative.

### Triplicate TMA cores provide adequate sampling for the classification of LMS as containing high or low TIL burden

To validate our finding from the vTMA experiment that a conventional number of TMA cores was sufficient for categorising tumours as having ‘high’ or ‘low’ TIL burden, we constructed a physical TMA (pTMA) that included triplicate 1mm cores from sampled tumours. 44/47 LMS were included in this TMA. In 11/44 (25%) tumours, the same block was used for pTMA construction as was used for the vTMA model. In 33/44 (75%) tumours, due to insufficient tissue depth remaining in blocks used for the vTMA, a different tumour block from the same specimen was used for core sampling for the pTMA.

In the vTMA model, a median of 11 cores were required to accurately estimate the absolute value for true TIL burden. We thus assessed if the triplicate cores used in pTMA were similarly inaccurate in estimating absolute true TIL burdens (defined as the mean of all 20 cores from the vTMA experiment as shown in **Figure 3C**) **(Figure 4A-C)**. Comparison of pTMA estimates to true TIL burdens using the Bland-Altman method for all 44 LMS cases indicated that the pTMA produced a modest overestimate of ‘true’ TIL burden (pTMA bias +46% for CD3, +9% for CD8), but that levels of agreement between pTMA-derived estimates and true TIL burdens were poor. The wide 95% limits of agreement detected in this analysis indicated that for any pTMA-derived estimate within the cohort, there would be 95% confidence that the associated ‘true’ TIL burden was anything from 6-8 times less or 9-14 times more than the estimate. These levels of agreement were improved when analysis was limited to the 11 tumours where pTMA and vTMA were taken from the same tumour blocks, but still reflected that estimates were associated with 95% confidence of true TIL burden falling between 2 times less or 3 times more the estimated value. These data show that pTMA estimates that are based on triplicate core sampling do not accurately estimate the true TIL burden, a finding that is consistent with results from the vTMA model.

**Figure 4.**
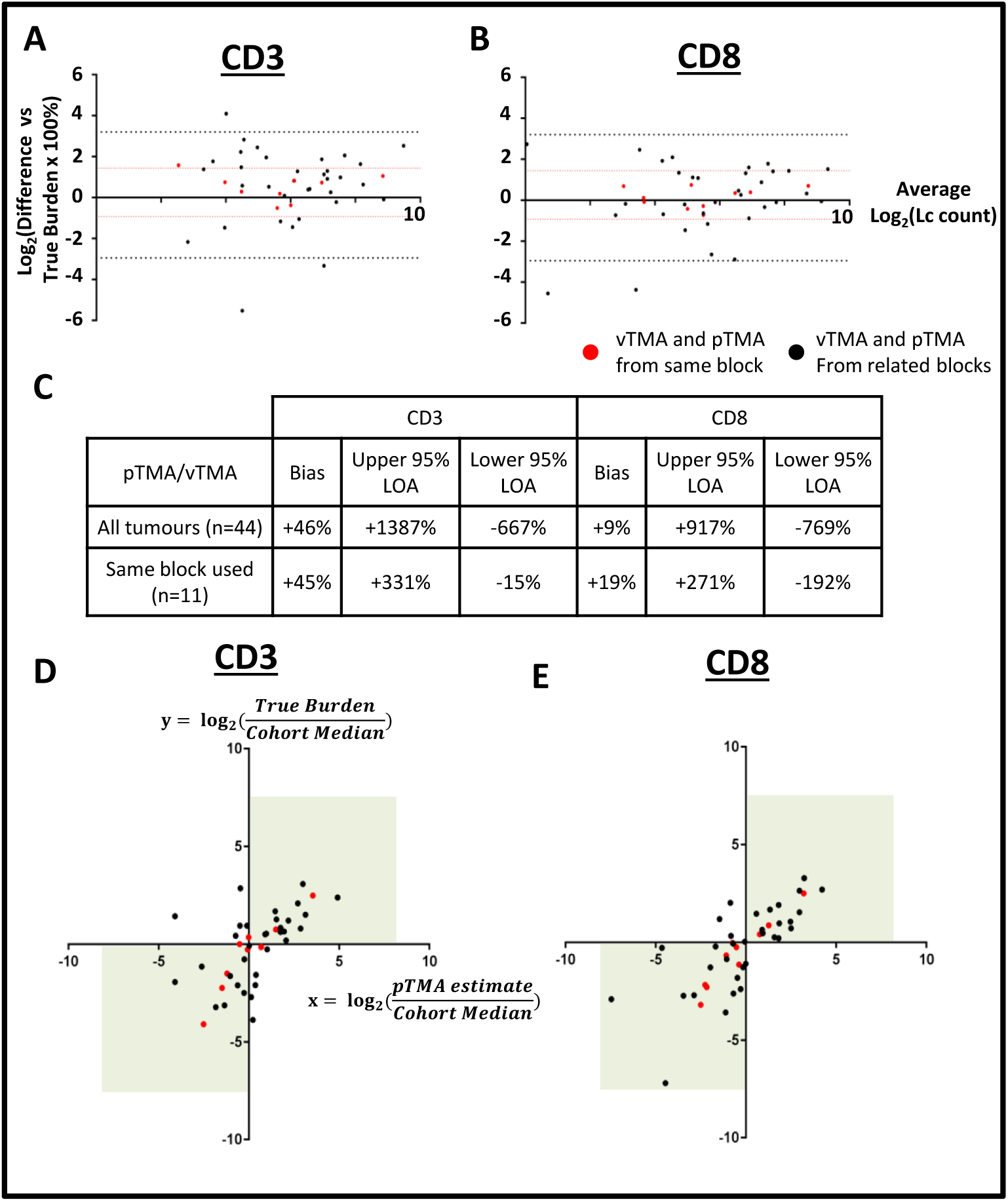
Triplicate TMA cores can identify tumours as having a high or low TIL burden, but do not accurately estimate precise TIL numbers. Bland Altman plots show percentage difference of pTMA-derived estimated TIL burdens compared to true TIL burdens for **(A)** CD3 and **(B)** CD8. 95% limits of agreement (LOA) for all 44 tumours shown by black dotted lines, 95% LOA for 11 tumours with pTMA and vTMA from same block shown by red dotted lines. LOA and biases from these plots are summarised in **(C).** Dot plots show ratio of pTMA-derived estimated TIL burden:cohort median value (x axis) plotted against ratio of true TIL burden:cohort median value (y axis) for **(D)** CD3+ and **(E)** CD8+ TILs. Ratio >0 indicates tumour identified as ‘High TIL burden’ (i.e above cohort median). Ratio <0 indicated tumour identified as ‘Low TIL burden’. Values in top right or bottom left quadrant (green boxes) indicate consistent TIL categorisation based on pTMA-derived estimate and true TIL burden. Red dots represent tumours where pTMA and vTMA sampled from same tumour block, black dots represent tumours where pTMA and vTMA were sampled from different blocks from same tumour specimen.

In the vTMA experiment, a median of 1 core was needed to correctly identify a LMS tumour has having a ‘high’ or ‘low’ CD3+ or CD8+ TIL burden, as defined by position above or below cohort median value. When triplicate cores within the pTMA were used to similarly assign tumours as having ‘high’ or ‘low’ TIL burdens (as per cohort median values, shown in **Figure 3C**), we found good levels of agreement with assignment versus the true TIL burden **(Figure 4D-E)**. Across all 44 tumours represented in the pTMA, accuracy (i.e. percentage of tumours that were correctly identified as having ‘high’ or ‘low’ true TIL burden) was 70.5% and 90.9% for CD3 and CD8 respectively. When limited to the 11 cases where the same block was used for both vTMA and pTMA, accuracy for correct identification of CD3+ or CD8+ TIL burden was improved to 72.7% and 100% respectively. Accuracy for the 33 cases where different blocks were used between vTMA and pTMA was 70.0% and 87.9%. These results demonstrate that, for a large majority of tumours in the cohort, triplicate TMA cores were adequate for correctly identifying whether the tumour had a ‘high’ or ‘low’ TIL burden. The accuracy of the pTMA was only modestly improved in tumours where the same tumour block was sampled for vTMA and pTMA, again indicating that there is only a minor contribution of intra-tumoural heterogeneity between related blocks to sampling error.

Consistent with the conclusions of the vTMA experiment (Figure 3D), these findings show that the inclusion of a conventionally used number of replicate cores from the same tumour (i.e. 3 cores) in a pTMA provides sufficient representation of true tumour TIL burden to accurately categorise tumours as having high or low TIL numbers, and that this accuracy is maintained between different blocks from the same tumour. However, the use of this relatively small number of cores can produce significant inaccuracy in estimating the absolute value of true TIL burden within a tumour.

## Discussion

In this study, we have characterised the TIL burden in a cohort of primary LMS tumours and demonstrate that there is evidence for both inter-and intra-tumoural heterogeneity in this STS subtype. We find that the TIL burden in LMS is generally low compared to immune-active cancer types such as melanoma and NSCLC, but that a subset of LMS exhibit heavier lymphocytic infiltration. Large intra-tumoural variation in TIL burden was observed in a minority of cases, particularly in tumours with a greater overall degree of TIL burden. However, across the whole cohort, the degree of intra-tumoural heterogeneity was small relative to the inter-tumoural differences in overall TIL burden between cases within the cohort. Additionally, our investigation of both a virtual and physical TMA indicates that a conventional and practical number of 1mm cores (between 1-3) provide sufficient representation for ordinal categorisation of tumours as having either a high or low degree of lymphocyte infiltration. These data indicate that intra-tumour heterogeneity of TIL burden may not be a great source of confounding sampling error and that TMAs represent a feasible and appropriate research tool for future immune profiling studies in LMS.

Our finding that TIL burdens are generally low in LMS is consistent with other studies that have used histological or gene expression deconvolution approaches to profile immune responses in LMS and other STS subtypes^25,38–40^. We also observe that a small number of LMS cases contain a higher degree of lymphocytic infiltrate and further studies in larger LMS cohorts are required to assess whether such differences in TIL burden can provide prognostic information or serve as predictive biomarkers for immunotherapies and/or other treatment modalities. Reported data have indicated that the biological and clinical relevance of TIL and other immune factors may vary between different STS subtypes^25,39–41^ – our focus on a single, more common STS subtype enables interpretation of our results without potential confounding by histological subtype-specific variation.

The applicability of our findings to other STS or epithelial cancers remains to be determined. Intra-tumoural heterogeneity in the immune microenvironment has been described in numerous epithelial cancer types, both within primary lesions and between different metastatic sites^7,11–15^. In breast cancer, a vTMA methodology was used to demonstrate that agreement between TMA and whole tumour assessment of TIL burden plateaued when sampling any more than four 0.6mm cores^23^. Interestingly, the degree of this correlation varied depending upon breast cancer subtype – Her2+ breast cancers had generally worse correlation, indicating greater spatial heterogeneity in TIL distribution – and that a greater degree of TIL ‘skewness’ (i.e. greater spatial heterogeneity) was itself independently associated with worse prognosis. This suggests that spatial uniformity of TIL burden may vary between different cancer types and within different molecular and histological subtypes, and that spatial distribution itself may provide clinically relevant information – both findings that warrant caution when adopting TMA methods for assessing TIL burden. Furthermore, while TILs are accepted as central mediators of anti-tumour immune responses, the immune-tumour microenvironment constitutes a broad and complex range of cellular and protein factors that actively determine the nature and clinical consequence of any tumour-related immune response^42^. Our study does not provide any direct information on the use of TMAs to assess non-TIL immune factors and the extrapolation of our findings beyond TILs remains to be investigated.

Our data indicate that TMAs can provide a degree of representation of overall tumour TIL burden which is adequate for ordinal categorisation into high or low subgroups. The design of future studies of the immune microenvironment of tumours should acknowledge the inherent limitations of TMA methods and consider the incorporation of additional orthogonal approaches such as gene expression analysis and flow cytometry methodologies that are capable of providing complementary information regarding the composition of immune subsets leading to a more comprehensive and accurate representation of tumour immune microenvironment.

## Acknowledgements

The authors acknowledge the support of The Royal Marsden Cancer Charity Sarcoma Research Fund and wish to thank the Breast Cancer Now Histopathology Core Facility for assistance in preparation of histology slides

This study represents independent research that was in part funded by a grant to IJ and RLJ from the National Institute for Health Research (NIHR) Biomedical Research Centre at The Royal Marsden. The views expressed are those of the author(s) and not necessarily those of the NHS, the NIHR or the Department of Health and Social Care.

## Author Contributions

ATJL, IJ, KT, PHH and RLJ conceived of and designed the study. MJS, DCS, AJH provided surgical specimens and technical annotation of tumour material. KT and CF performed diagnostic assessment and annotation of tumour material. ATJL and KT performed histological assessment and data collection. ATJL and WC performed statistical analysis and figure preparation. AL, WC, RLJ and PHH prepared initial manuscript drafts which were then reviewed by all authors.

## Additional information

All authors declare no competing interests

## References

1 Kononen, J. et al. Tissue microarrays for high-throughput molecular profiling of tumor specimens. Nat. Med. 4, 844–8471 (1998).

2 Bedard, P. L., Hansen, A. R., Ratain, M. J. & Siu, L. L. Tumour heterogeneity in the clinic. Nature 501, 355–364 (2013).

3 Camp, R. L., Neumeister, V. & Rimm, D. L. A decade of tissue microarrays: Progress in the discovery and validation of cancer biomarkers. J. Clin. Oncol. 26, 5630–5637 (2008).

4 Kallioniemi, O. P., Wagner, U., Kononen, J. & Sauter, G. Tissue microarray technology for high-throughput molecular profiling of cancer. Hum. Mol. Genet. 10, 657–662 (2001).

5 Camp, R. L., Charette, L. a & Rimm, D. L. Validation of tissue microarray technology in breast carcinoma. Lab. Invest. 80, 1943–1949 (2000).

6 Rubin, M. A., Dunn, R., Strawderman, M. & Pienta, K. J. Tissue microarray sampling strategy for prostate cancer biomarker analysis. Am. J. Surg. Pathol. 26, 312–9 (2002).

7 Galon, J. et al. Type, Density, and Location of Immune Cells Within Human Colorectal Tumors Predict Clinical Outcome. 313, 1960–1965 (2006).

8 Denkert, C. et al. Tumor-Infiltrating Lymphocytes and Response to Neoadjuvant Chemotherapy With or Without Carboplatin in Human Epidermal Growth Factor Receptor 2-Positive and Triple-Negative Primary Breast Cancers. J. Clin. Oncol. 33, 983–991 (2015).

9 Galon, J. et al. Immunoscore and Immunoprofiling in cancer: an update from the melanoma and immunotherapy bridge 2015. J. Transl. Med. 14, 273 (2016).

10 Herbst, R. S. et al. Predictive correlates of response to the anti-PD-L1 antibody MPDL3280A in cancer patients. Nature 515, 563–567 (2014).

11 Kaur, H. B. et al. Association of tumor-infiltrating T-cell density with molecular subtype, racial ancestry and clinical outcomes in prostate cancer. Mod. Pathol. (2018). doi:10.1038/s41379-018-0083-x

12 Baras, A. S. et al. The ratio of CD8 to Treg tumor-infiltrating lymphocytes is associated with response to cisplatin-based neoadjuvant chemotherapy in patients with muscle invasive urothelial carcinoma of the bladder. Oncoimmunology 5, e1134412 (2016).

13 Wei, J. S. et al. Clinically Relevant Cytotoxic Immune Cell Signatures and Clonal Expansion of T Cell Receptors in High-risk MYCN -not-amplified Human Neuroblastoma. Clin. Cancer Res. clincanres.0599.2018 (2018). doi:10.1158/1078-0432.CCR-18-0599

14 Kluger, H. M. et al. Characterization of PD-L1 Expression and Associated T-cell Infiltrates in Metastatic Melanoma Samples from Variable Anatomic Sites. Clin. Cancer Res. 21, 3052–60 (2015).

15 Schalper, K. A. et al. Objective Measurement and Clinical Significance of TILs in Non–Small Cell Lung Cancer. JNCI J Natl Cancer Inst 107, (2015).

16 Jiménez-Sánchez, A. et al. Heterogeneous Tumor-Immune Microenvironments among Differentially Growing Metastases in an Ovarian Cancer Patient. Cell 170, 927–938.e20 (2017).

17 Shi, L. et al. Multi-omics study revealing the complexity and spatial heterogeneity of tumor-infiltrating lymphocytes in primary liver carcinoma. Oncotarget 8, 34844–34857 (2017).

18 Li, M. et al. Heterogeneity of PD-L1 expression in primary tumors and paired lymph node metastases of triple negative breast cancer. BMC Cancer 18, 1–9 (2018).

19 Rehman, J. A. et al. Quantitative and pathologist-read comparison of the heterogeneity of programmed death-ligand 1 (PD-L1) expression in non-small cell lung cancer. Mod. Pathol. 30, 340–349 (2017).

20 Mlecnik, B. et al. Comprehensive Intrametastatic Immune Quantification and Major Impact of Immunoscore on Survival. JNCI J. Natl. Cancer Inst. 110, 97–108 (2018).

21 Zlobec, I., Koelzer, V. H., Dawson, H., Perren, A. & Lugli, A. Next-generation tissue microarray (ngTMA) increases the quality of biomarker studies: an example using CD3, CD8, and CD45RO in the tumor microenvironment of six different solid tumor types. J. Transl. Med. 11, 104 (2013).

22 Botti, G., Scognamiglio, G. & Cantile, M. PD-L1 immunohistochemical detection in tumor cells and tumor microenvironment: Main considerations on the use of tissue micro arrays. Int. J. Mol. Sci. 17, 8–10 (2016).

23 Khan, A. M. & Yuan, Y. Biopsy variability of lymphocytic infiltration in breast cancer subtypes and the ImmunoSkew score. Sci. Rep. 6, 2–11 (2016).

24 George, S., Serrano, C., Hensley, M. L. & Ray-Coquard, I. Soft Tissue and Uterine Leiomyosarcoma. J. Clin. Oncol. 36, 144–150 (2018).

25 Abeshouse, A. et al. Comprehensive and Integrated Genomic Characterization of Adult Soft Tissue Sarcomas. Cell 171, 950–965.e28 (2017).

26 Davoli, T., Uno, H., Wooten, E. C. & Elledge, S. J. Tumor aneuploidy correlates with markers of immune evasion and with reduced response to immunotherapy. Science (80-.). 355, eaaf8399 (2017).

27 Patnaik, A. et al. Phase I Study of Pembrolizumab (MK-3475; Anti-PD-1 Monoclonal Antibody) in Patients with Advanced Solid Tumors. Clin. Cancer Res. 21, 4286–93 (2015).

28 George, S. et al. Loss of PTEN Is Associated with Resistance to Anti-PD-1 Checkpoint Blockade Therapy in Metastatic Uterine Leiomyosarcoma. Immunity 46, 197–204 (2017).

29 D’Angelo, S. P. et al. Nivolumab with or without ipilimumab treatment for metastatic sarcoma (Alliance A091401): two open-label, non-comparative, randomised, phase 2 trials. Lancet Oncol. 19, 416–426 (2018).

30 Fletcher, C. D. M., Hogendoorn, P. C. W., Mertens, F. & Bridge, J. A. WHO Classification of Tumours of Soft Tissue and Bone. (IARC Press, 2013).

31 Schindelin, J. et al. Fiji: an open-source platform for biological-image analysis. Nat. Methods 9, 676–682 (2012).

32 Wolak, M. E., Fairbairn, D. J. & Paulsen, Y. R. Guidelines for estimating repeatability. Methods Ecol. Evol. 3, 129–137 (2012).

33 Anitei, M.-G. et al. Prognostic and Predictive Values of the Immunoscore in Patients with Rectal Cancer. Clin. Cancer Res. 20, 1891–1899 (2014).

34 Pagès, F. et al. International validation of the consensus Immunoscore for the classification of colon cancer: a prognostic and accuracy study. Lancet 391, 2128–2139 (2018).

35 Rao, U. N. M. et al. Presence of tumor-infiltrating lymphocytes and a dominant nodule within primary melanoma are prognostic factors for relapse-free survival of patients with thick (t4) primary melanoma: pathologic analysis of the e1690 and e1694 intergroup trials. Am. J. Clin. Pathol. 133, 646–53 (2010).

36 Geng, Y. et al. Prognostic Role of Tumor-Infiltrating Lymphocytes in Lung Cancer: a Meta-Analysis. Cell. Physiol. Biochem. 37, 1560–1571 (2015).

37 Hendry, S. et al. Assessing Tumor-Infiltrating Lymphocytes in Solid Tumors. Adv. Anat. Pathol. 24, 311–335 (2017).

38 Thorsson, V. et al. The Immune Landscape of Cancer. Immunity 48, 812–830.e14 (2018).

39 D’Angelo, S. P. et al. Prevalence of tumor-infiltrating lymphocytes and PD-L1 expression in the soft tissue sarcoma microenvironment. Hum. Pathol. 46, 357–65 (2015).

40 Sorbye, S. W. et al. Prognostic Impact of Lymphocytes in Soft Tissue Sarcomas. PLoS One 6, e14611 (2011).

41 Boxberg, M. et al. PD-L1 and PD-1 and characterization of tumor-infiltrating lymphocytes in high grade sarcomas of soft tissue–prognostic implications and rationale for immunotherapy. Oncoimmunology 7, (2018).

42 Chen, D. S. et al. Oncology Meets Immunology: The Cancer-Immunity Cycle. Immunity 39, 1–10 (2013).

